# Improved Serodiagnosis of *Trypanosoma vivax* Infections in Cattle Reveals Higher Infection Rates in the Livestock Regions of Argentina

**DOI:** 10.1101/2024.02.22.581515

**Authors:** Iván Bontempi, Diego G. Arias, Graciela V. Castro, Luz Peverengo, Genaro Díaz, Martín Allassia, Gonzalo Greif, Iván Marcipar

## Abstract

Bovine trypanosomiasis, caused by Trypanosoma vivax, currently affects cattle, resulting in significant economic consequences in sub-Saharan Africa and South America. The development of new diagnostic antigens is crucial for improving and refining existing methods. Our study assessed the effectiveness of two recombinant antigens in detecting specific antibodies in cattle. These antigens are derivatives of an invariant surface glycoprotein (ISG) from T. vivax. We evaluated a fraction of an antigen previously described (TvY486_0045500), referred to as TvISGAf, from an African strain, and identified a new ISG antigen from an American isolate, TvISGAm. ELISA evaluation using these antigens was conducted on 212 samples from cattle. The diagnostic performance was enhanced when utilizing a combination of both antigens (denominated TvISG-based ELISA), achieving a sensitivity of 89.6% and specificity of 93.8%. Following validation of the TvISG-based ELISA, we determined the seroprevalence of T. vivax infection in 892 field samples from cattle in the central region of Argentina. The average seroprevalence of T. vivax was 53%, with variation across the six surveyed departments ranging from 21% to 69%. These results support the use of the TvISG ELISA as a valuable serological tool for detecting and monitoring T. vivax infection in cattle. They also reported for the first time T. vivax seroprevalence in Argentina, highlighting the widespread endemic nature of the disease in the region. To effectively manage the increasing spread of T. vivax in the vast livestock production areas of South America, we support the need for consistent surveillance programs and implementation of preventive strategies.

## Introduction

African Animal Trypanosomiasis (AAT), caused by *Trypanosoma vivax (T. vivax)*, is a disease that threatens the health and productivity of cattle in Africa and Latin America (1). In South America, *T*. *vivax* is the primary causative agent of AAT and is mainly transmitted mechanically by hematophagous flies of the Stomoxys and Tabanus genera owing to the absence of a biological vector, the tsetse fly (2). In this region, AAT poses a potential risk to nearly 350 million cattle, with studies on outbreaks in the Pantanal region of Brazil and Bolivia estimating a potential loss of US$160 million (3).

Among the trypanosomes transmitted by the tsetse fly (Salivaria), *T. vivax* is phylogenetically positioned in the early branch (4,5) and belongs to the subgenus Duttonella, which predominantly causes severe wasting disease in cattle but may also infect sheep, goats, camels, horses, and buffalo (1,6). Acute *T. vivax* infection is characterized by anemia, fever, poor body condition, and abortions. Occasionally, the disease may progress to more severe disorders, such as severe neurological symptoms, and can be fatal (7–9). Chronic *T. vivax* infection leads to progressive anemia, weight loss, and reproductive failure, resulting in a noticeable reduction in milk production (10,11). However, infected cattle can be asymptomatic (12), and if the infection is left untreated, the animals can become asymptomatic carriers, potentially spreading and perpetuating the disease throughout the herd (13). Although the earliest reports in South America date back to French Guiana in 1919 (14), the first outbreak recorded in Argentina was in 2006 in Formosa, northeastern Argentina (15). Argentina is one of the world’s leading beef exporters, with northeastern and central-eastern areas being vital livestock production zones. Despite initial reports in Argentina, *T. vivax* was recently identified through molecular methods in 2018 (16) and has been associated with acute disease in dairy cattle in the Pampas region, having a substantial economic impact (17).

Diagnosis of *T. vivax* trypanosomiasis must rely on highly sensitive and specific methods capable of detecting cryptically infected animals that might serve as healthy carriers, while also distinguishing between *T. vivax* and other trypanosomes, such as *Trypanosoma theileri* and *Trypanosoma evansi*. Molecular assays; such as conventional PCR; have successfully detected *T. vivax* (4,11,17). However, these assays are costly and require specialized laboratories for execution, rendering them inaccessible to field veterinarians and dairy farm owners. The use of serological techniques has come into play, enabling the assessment of Trypanosoma presence in a large sample volume at an affordable cost. Serological diagnostic techniques for AAT are based on the detection of antibodies against parasites using an indirect fluorescent antibody test (IFAT), as described by Luckins and Mehlitz in 1978 (18). Although IFAT is sensitive and specific, it is not quantitative, requires fluorescence-enabled microscopes, and lacks standardized antigen preparation. Whole trypanosomal lysate was also used for the diagnosis of AAT through ELISA (19). Although it exhibits high sensitivity and specificity, it also has the drawback of requiring constant parasite production, making its preparation and standardization challenging. Assays employing recombinant proteins have eliminated the need for live parasite preparation. Different antigens have been proposed for the diagnosis of bovine trypanosomiasis, including MyxoTLm (20), GM6 (21,22), variant surface glycoproteins (VSG) (23) and invariant surface glycoproteins (ISGs) (24,25).

ISGs are conserved antigens with unknown functions, are expressed at relatively low levels on the trypanosome membrane, and exhibit no known antigenic variation (26). In recent years, ISGs have gained attention for their conserved functionality in diagnostic roles, detecting *Trypanosoma brucei gambiense* in Human African Trypanosomiasis (27), and for both *Trypanosoma congolense* and *T. vivax* in AAT (24,25). Fleming et al. (2016) identified an ISG with high diagnostic potential, called TvY486_0045500. However, it was amplified from an African strain and evaluated using sera from African cattle infected with *T. vivax*. In the present study, we developed a diagnostic method employing the TvY486_0045500 antigen and ISG from an American *T. vivax* isolate. We assessed its performance using sera from cattle infected with the American *T. vivax* strains. In addition, using the developed assay, we determined the prevalence of *T. vivax* in the dairy basin of Argentina.

## Methods

### Materials

The bacteriological medium was purchased from Britania Laboratories. *Taq* and *Pfu* DNA polymerase, T4 DNA ligase, and restriction enzymes were purchased from Promega and Thermo Fisher Scientific. Amylose resin was purchased from New England Biolabs. Ni^2^-HiTrapTM chelating HP column was purchased from GE Healthcare. All other reagents and chemicals were of the highest quality and were commercially available from Sigma-Aldrich and Merck.

### Bacteria, plasmids, genetic material and *T. vivax* cells

*Escherichia coli* TOP10 F’ (Invitrogen) and *E. coli* BL21 (DE3) (Novagen) cells were used for routine plasmid construction and expression experiments, respectively. The pGEM-T Easy vector (Promega) was used for cloning and sequencing. The expression vectors used were pET28 (for N– and C-terminal His-tag protein generation, Novagen) and pMOMAL (a homemade-derived plasmid from pMAL C2 with recombinant His-tags and MBP-tags). Genomic DNA from *T. vivax* was obtained using the Wizard Genomic DNA Purification Kit (Promega). DNA manipulation, *E. coli* culture, and transformation were performed according to the standard protocols (28).

### Molecular cloning

Based on the available information on *T. vivax* genome sequences (https://tritrypdb.org/), ISG-encoding sequences were amplified by PCR using genomic DNA from the African strain of *T. vivax* Y486 (TvY486_0045500 – https://tritrypdb.org/tritrypdb/app) and *T. vivax* isolated in Argentina (tig00000163 –(29)). PCR was performed using genomic DNA and specific primer pairs (S1 Table) under the following conditions: 95 °C for 10 min; 30 cycles of 95 °C for 30 s, 55-65 °C for 30 s, 72 °C for 1.5 min, and 72 °C for 10 min. The PCR product was subsequently purified and ligated into the pGEM-T Easy vector and its fidelity and identity were confirmed by complete DNA sequencing (Macrogen, South Korea). The obtained pGEM-T Easy constructs were digested with the appropriate restriction enzymes and the purified amplicons were ligated to different expression vectors using T4 DNA ligase (Promega) for 16 h at 4 °C. Truncated versions of the African and American ISG sequences, TvISGAf and TvISGAm, were cloned into the pMOMAL and pET28 plasmids, respectively. Competent *E. coli* BL21 (DE3) cells transformed with the construct were selected in agar plates containing Lysogeny Broth (LB; 10 g L^-1^ NaCl, 5 g L^-1^ yeast extract, 10 g L^-1^ peptone, pH 7.4) supplemented with ampicillin (100 μg mL^-1^) or kanamycin (50 μg mL^-1^) as appropriate. Plasmid DNA was prepared and subsequent restriction treatment was performed to check the correctness of the construct.

### Overexpression and purification of the recombinant protein

A single colony of *E. coli* BL21 (DE3), transformed with each recombinant plasmid, was selected. Overnight cultures were diluted 1/100 in fresh LB medium supplemented with the appropriate antibiotic and grown under identical conditions to the exponential phase, with an OD_600_ of 0.6. Expression of the respective recombinant proteins was performed with 0.25 mM IPTG for 16 h at 23 °C. The cells were harvested and stored at –20 °C until further use.

Purification of each recombinant protein was performed using IMAC with a 1 mL Ni^2+^-HiTrap^TM^ chelating HP column (GE Healthcare). Briefly, the bacterial pellet was resuspended in binding buffer (20 mM Tris–HCl pH 7.5, 400 mM NaCl, and 10 mM imidazole) and disrupted by sonication using a high-intensity ultrasonic processor (Vibra-cell TM VCX-600; Sonics & Materials Inc.). The lysate was centrifuged (10000 ×*g* for 30 min) to remove cell debris. The resultant crude extract was loaded onto a column that had been equilibrated with binding buffer. After washing with 10 bead volumes of the same buffer, the recombinant protein was eluted with elution buffer (20 mM Tris–HCl pH 7.5, 400 mM NaCl, 300 mM imidazole). In addition, isolated TvISGAf from IMAC chromatography (as an MBP-fusion protein) was purified using amylose affinity chromatography (using a 5 mL amylose column, New England Biolabs), as a complementary step, according to the manufacturer’s instructions. Fractions containing pure proteins were pooled, concentrated, dialyzed (using 20 mM Tris-HCl buffer pH 8.0, 200 mM NaCl, and 1 mM EDTA), and frozen with 20% (v/v) glycerol at –80 °C.

### Protein methods and immunodetection

Protein concentration was determined following the procedure described by Bradford (30) using bovine serum albumin (BSA) as a standard.

Immunodetection experiments were performed using *T. vivax* cells from Y486 isolated from the blood of infected mice for four days. This protocol was adapted from a previously described method for *T. cruzi* (31). Polyclonal antibodies against TvISGAf or TvISGAm were produced by immunizing mice with purified recombinant antigens as previously described (32). Mouse anti-TvISGAf or anti-TvISGAm polyclonal antibodies, rabbit anti-*Tc*cPx polyclonal antibodies (1/100 dilution), and FITC-conjugated goat anti-rabbit or Alexa 680-conjugated goat anti-mouse antibodies (1/1000 dilution) were used for protein labeling. The slides were mounted in the presence of anti-fade mounting solution (supplemented with DAPI) and visualized under a confocal microscope (Leica).

### Parasitological diagnosis

For *Trypanosoma* spp. diagnosis, blood samples were promptly transferred into glass microhematocrit capillary tubes containing sodium heparin (80 U/mL, Biocap) within 6 hours of collection. The capillary tubes were then centrifuged in a microhematocrit centrifuge at 9000 rpm for 5 min. Subsequently, packed cell volumes were determined. Trypanosome motility was assessed according to previously established protocols (33). A molecular diagnosis was made to confirm infections with *T. vivax*. Blood was collected from the jugular veins of dairy cattle by using guanidine/EDTA and stored at 4 °C until further use. Genomic DNA was extracted from 200 µL of blood using the High Pure PCR Template Preparation Kit (Roche), following the manufacturer’s instructions. PCR amplification targeting sequences corresponding to the catalytic domain of CatL-like (cdCatL-like) enzymes was performed using the oligonucleotide primers TviCatL1 and DTO155 (4). PCR assays were conducted in 25 µL reaction volumes comprising 1X reaction buffer (100 mM Tris-HCl, 50 mM KCl, pH 8.8), 0.2 mM dNTP, 1–4 mM MgCl_2_, 0.4 µM of each primer, 0.625 U Taq polymerase (Promega), and 5 µL of DNA samples. The PCR products were visualized using 2% agarose gel electrophoresis, stained with GelRed™ (Millipore), and observed under UV light. Furthermore, a 383-bp fragment from the Cyt b gene present in mammals was amplified for all samples that yielded negative results using the molecular method (34). This was performed to assess the DNA integrity and quality, including the presence of inhibitors. Samples that failed to amplify were excluded from subsequent data analysis.

### Positive and Negative Serum Samples

The serum panel consisted of 212 serum samples, 149 positive and 63 negative, from dairy cattle, mainly the Holstein and Jersey breeds. Positive serum samples were collected on either day 15 or 21 after confirmation by microscopic examination of thin blood smears, buffy coat, and TviCatL-PCR

(4). For the screening evaluation, we selected 30 positive and 30 negative samples from the entire panel. Finally, serum samples from cattle infected with *Babesia bovis* (n = 20*), Anaplasma marginale* (n = 20), and *Trypanosoma theileri* (n = 8) were used as negative controls for infections caused by unrelated hemoprotozoan parasites. These infections were confirmed by observation of parasitemia and detection of specific antibodies in cattle blood and sera.

For kinetic evaluation, 50 positive serum samples were collected from Holstein dairy cattle on day 0 after confirmation by microscopic examination of thin blood smears, buffy coats, and TviCatL-PCR. Following the confirmation of the presence of *T. vivax*, the bovines were treated with isometamidium (ISM, OVER). Samples from the same Holstein dairy cattle were collected 15 and 30 days after the initial detection of *T. vivax*. Among these, 14 of the 50 positive serum samples originated from a dairy farm with a recent *T. vivax* infection report and were denoted as the acute group. The remaining 36 of the 50 positive serum samples belonged to a dairy farm with a history of chronic *T. vivax* infection, termed the chronic group.

### Field Survey

A comprehensive field survey was conducted in the provinces of Santa Fe and Córdoba, in the central region of Argentina. In these provinces, the sampled animals primarily consisted of dairy cows (mainly Holstein and Jersey cows). This region witnessed numerous severe outbreaks of *T. vivax* infection in 2017, and the disease remains endemic in this area (17). Six areas were strategically selected from the regions with substantial livestock populations. The geographical locations and names of the six surveyed areas, all situated within the central region, are shown on a map (Fig. 1). The infected animals were routinely treated with diminazene acetate. Blood samples were collected between January 2021 and January 2023. Approximately 5 mL of blood was extracted from the jugular vein and placed in plastic tubes for serum preparation to ensure ambient temperature conditions. Subsequently, the serum samples were stored at −20 °C. A total of 892 blood samples were randomly obtained from cattle across a span of two years (2021–2023) from a total of 40 small farms. These samples were then subjected to examination using the microhematocrit technique (MHT), as previously described (Woo, 1969), within 3–6 h after blood collection. Each animal was sampled only once during the course of the study, using a cross-sectional approach. Microscopic analysis of Giemsa-stained blood smears from the afflicted animals was performed to assess the presence of bovine babesiosis and anaplasmosis.

**Figure 1:**
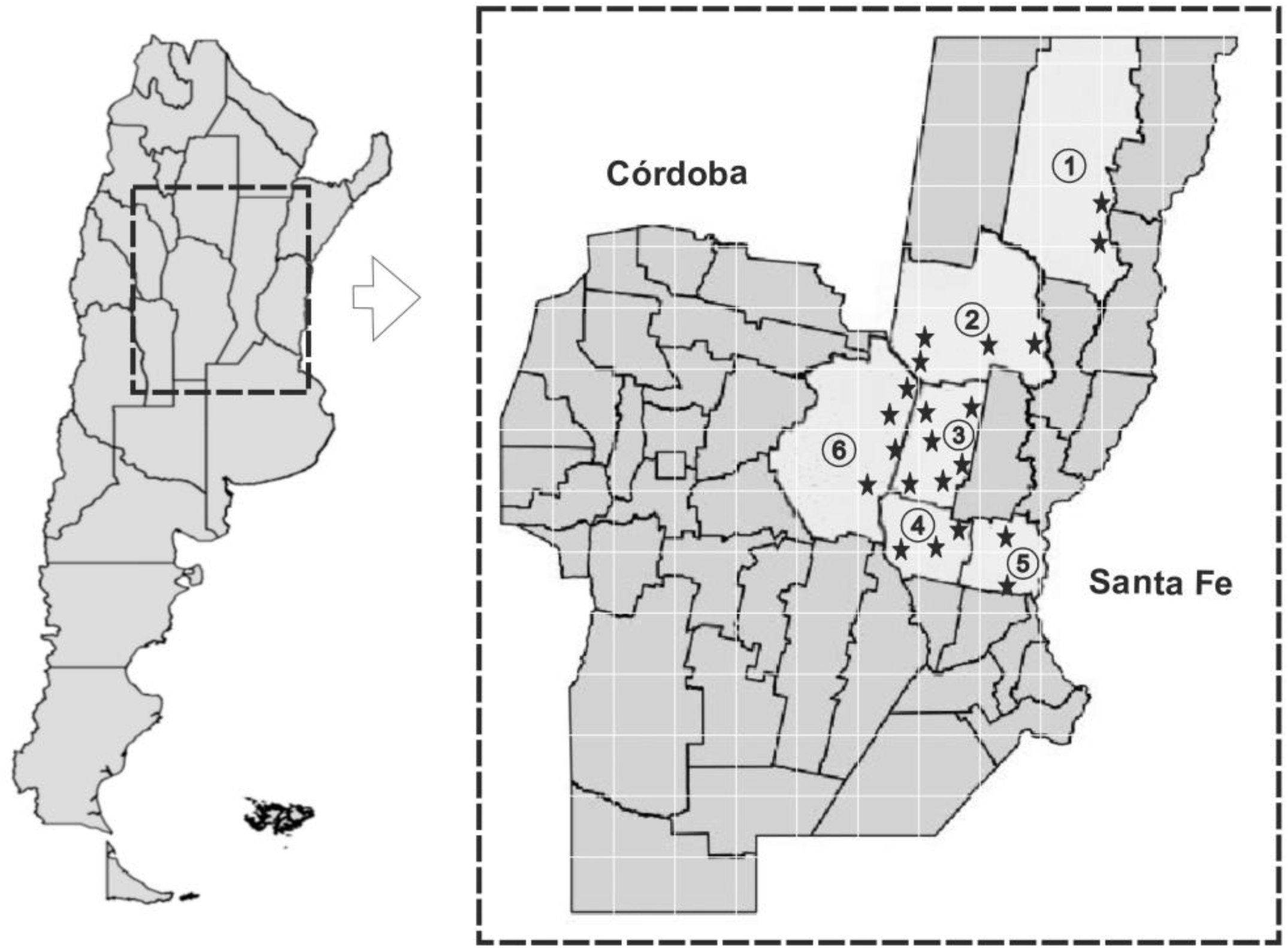
Map of Argentina showing the location of the two provinces surveyed in the study during 2021–2023. Identification Number (IN) indicated departments: ①: Vera, ②: San Cristóbal, ③: Castellanos, ④: San Martin, ⑤: San Jeronimo, ⑥: San Justo. **Star** indicate cities, towns, villages, communes.

### ELISA assay

The indirect ELISA was performed according to the current protocol. Microtiter plates (Greiner) were coated with 500 ng of specific antigens (TvISGAf and/or TvISGAm) in 50 mM carbonate-bicarbonate buffer (pH 9.6) and incubated overnight at 4 °C. The plates were washed thrice with 0.01% Tween 20 in phosphate-buffered saline (PBS-T) and then blocked with PBS containing 5% skimmed milk. Following another wash with PBS-T, serum samples were added to each well (diluted 40 times in 1% skimmed milk in PBS) and incubated at 37 °C for 1 h. A subsequent round of washing was performed, and peroxidase-conjugated goat anti-bovine IgG (Sigma-Aldrich) was added and incubated at 37 °C for 1 h. For substrate application, 3, 3’, 5, 5’;-tetramethylbenzidine (Invitrogen™) was used, and the reaction was halted by the addition of 1 M sulfuric acid. The optical density (OD) was measured at 450 nm using an ELx808 microplate reader (BioTek). For each specific antigen, the results were presented as the average OD derived from two concurrent evaluations of the same serum sample. Within each plate and for each specific antigen, six negative controls (seronegative for *T. vivax*) were examined simultaneously. The cut-off values for the ELISA results were determined by calculating the mean OD of the negative control serum samples, in addition to two standard deviations. Antibody levels were quantified as the ratio between the OD of the sample and the cut-off value. This measure is referred to as IODN (index of the optical density of the antibodies relative to the negative control) (35). The IODN value below which it was considered negative was determined using ROC curve analysis.

## Data analysis

A Kolmogorov-Smirnov test was initially conducted to assess the normality of the samples prior to comparing positive and negative serum outcomes. If the samples exhibited a normal distribution, comparisons between groups were performed using unpaired Student’s t-test. For samples that did not display a normal distribution, the Mann-Whitney U test was used for comparisons. Statistical significance was set at *P* ≤ 0.05. The performance of each assay was assessed using sensitivity (Se), specificity (Sp), positive predictive value (PPV), negative predictive value (NPV), area under the curve (AUC), and accuracy (AC). ROC analysis was employed to determine the lower limit of positivity (cutoff) and ascertain the optimal combination of sensitivity, specificity, and AUC. The quality of each test was classified based on the AUC results from the ROC analysis as follows: “Excellent” (1.0– 0.9), “Good” (0.9–0.8), “Reasonable” (0.8–0.7), “Poor” (0.7–0.6), and “Fail” (< 0.6). To assess the relationship between variables (time vs. antibody levels (IODN)) within each group, Pearson’s correlation test was performed. Subsequently, a general linear model was applied to each group using regression analysis. For statistical analysis, GraphPad Prism 7.0 and MedCalc 12.2.1. software were used.

## Results

### *In silico* analysis of sequence encoding for a *T. vivax* invariant surface glycoprotein and its recombinant expression

The information available in the database of the *Trypanosoma vivax* Y486 genome project (http://tritrypdb.org/tritrypdb/) presents a 1203 bp open reading frame (TvY486_0045500) that encodes for an invariant surface glycoprotein (of 400 amino acids with a theoretical molecular mass of 44.5 kDa, S1 Fig, previously analyzed by Fleming et al., 2016. Based on this information, we searched the genomic data of an American *T. vivax* isolate (29) for a putative gene encoding an invariant surface glycoprotein. Using the BLASTn tool and nucleotide sequence of TvY486_0045500, we identified a nucleotide sequence encoding a putative invariant surface glycoprotein (tig00000163, (29)). The amino acid sequence is presented in S2 Fig. *In silico* analysis predicted that both invariant surface glycoproteins from African and American *T. vivax* exhibit a typical ISG domain structure with an N-terminal signal peptide, transmembrane domain, and intracellular domain (S1 and S2 Figs). Amino acid sequence analysis with African and American *T. vivax* ISG showed that these proteins exhibited ∼63% identity and ∼69% similarity (S3 Fig).

Based on previous information (24) and prediction of linear epitopes (S4 Fig), we amplified truncated ISG sequences from African (TvISGAf) and American (TvISGAm) *T. vivax* genomic DNA by PCR (without the signal peptide and N-terminal transmembrane domain). The resulting DNA sequences were cloned into a pGEM-T Easy vector for analysis. To characterize the antigenic functionality of the recombinant proteins, we expressed the truncated forms in bacteria as a fusion protein with an N-terminal MBP-His-tag (TvISGAf from AA 126-400) and an N– and C-double His-tag (TvISGAm from AA 125 to 274) for chromatographic purification (the amino acid sequences of the resultant recombinant proteins are shown in S5 Fig). Under the recombinant expression conditions, we detected the expression of truncated proteins in the soluble extracts of recombinant *E. coli* (data not shown). SDS-PAGE analysis showed that recombinant proteins were produced and isolated with high electrophoretic purity (Fig. 2). The purified recombinant proteins were stored at –80 °C for at least eight months.

**Figure 2:**
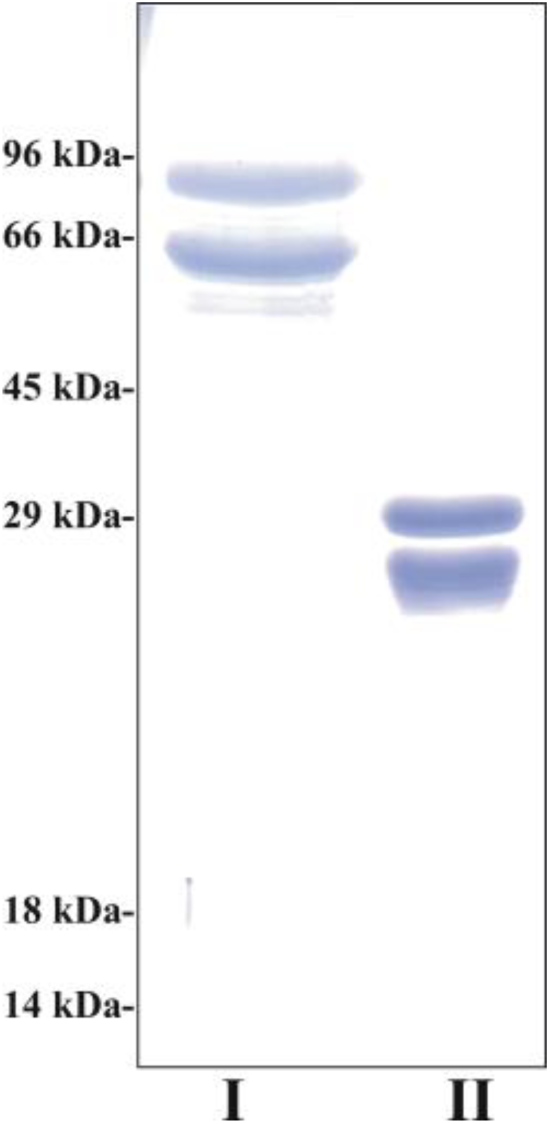
SDS-PAGE of purified recombinant proteins. Purified recombinant proteins were defined by electrophoretic migration under reducing and denaturing conditions, and subsequent staining with Coomassie blue. Lane I: TvISGAf (1 μg); Lane II: TvISGAm (1 μg).

As mentioned above, the SOSUI predictor server suggested that the ISG protein is a transmembrane protein. Using mouse-specific antibodies against recombinant TvISGAf and TvISGAm, we evaluated the localization of the ISG protein in African *T. vivax* cells. Using immunofluorescence microscopy (Fig. 3), we detected recognition signals throughout the parasite plasma membrane. This is in contrast to the signal observed for cytoplasmic peroxiredoxin, which presents a recognition signal throughout the cell (Fig. 3). These results support that ISG exhibit location on the cell surface in *T. vivax* cells.

**Figure 3:**
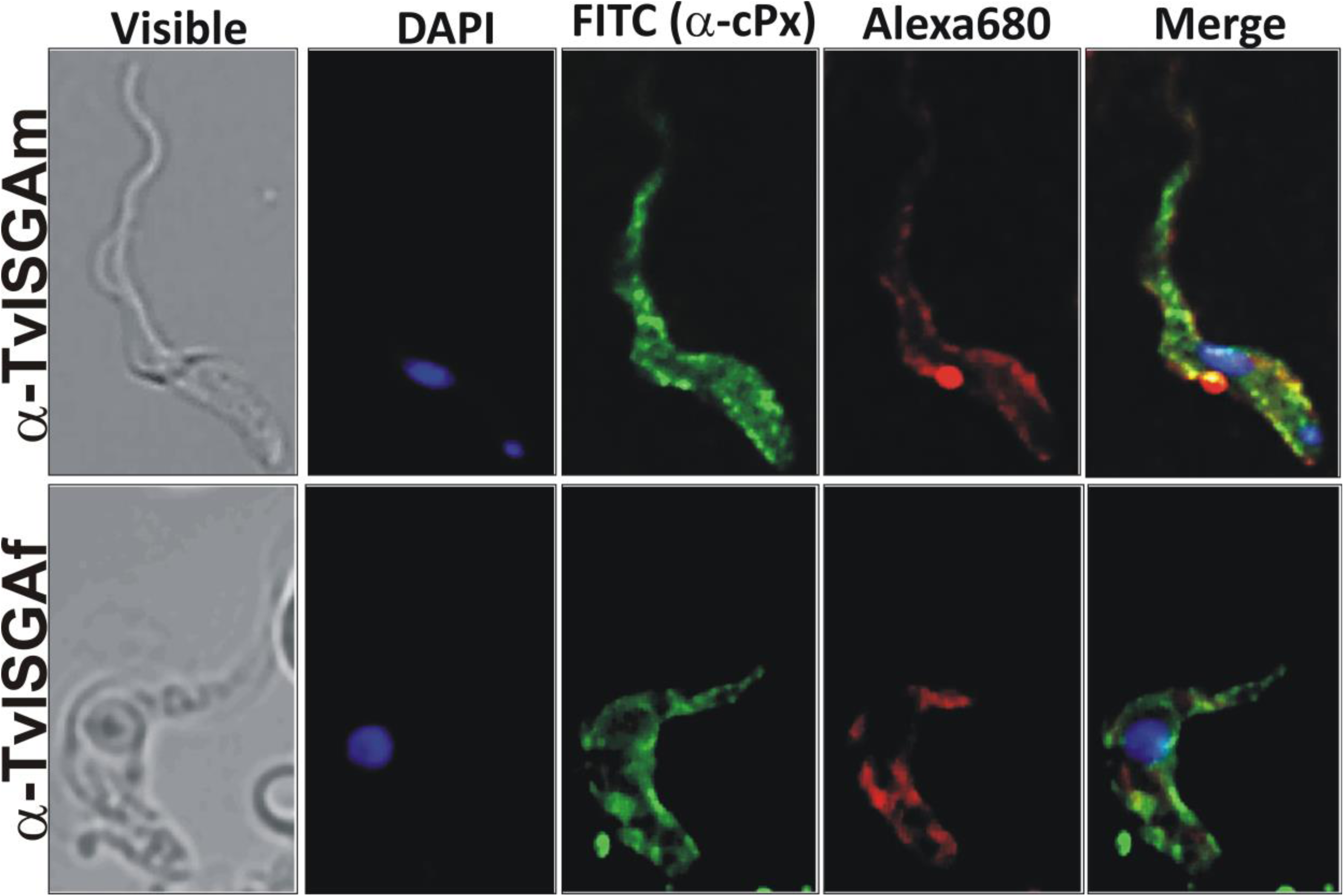
Immunodetection of ISG proteins in *T. vivax*. Confocal microscopy images was performed by the co-immunolocalization of ISG (labeled with mouse anti-TvISGAf or anti-TvISGAm antibodies and Alexa680 conjugated goat anti-IgG as a secondary antibody, red) and cytoplasmic peroxiredoxin (cPx, labeled with rabbit anti-cPx antibody and FITC-conjugated goat anti-IgG as a secondary antibody, green). DAPI was used for the nuclear and kinetoplast staining (blue). Negative controls (incubations without primary antibodies) are presented in the Supplementary Fig. S6.

### Preliminary ELISA screenings

The purified recombinant trypanosome proteins were used to prepare ELISA plates, as outlined in the Experimental Procedures section, and were examined against 60 samples of the sera panel, categorized into cattle infected with *T. vivax* (n = 30) and non-infected cattle (n = 30). Protein ELISA plates were assessed using all individual sera. These antigens encompassing TvISGAf (TvISGAf-based ELISA), TvISGAm (TvISGAm-based ELISA), and a combination of both proteins (TvISG-based ELISA) were evaluated. Diverse mixing conditions were evaluated for the proportion of each protein in the mixture, with a 1:1 proportion yielding optimal discrimination in the evaluated positive and negative pools (data not shown).

The data are presented as box plots for each distinct recombinant antigen ELISA plate (Fig. 4), offering visualization of the IODN range. The data indicated that both ELISA assays, one based on TvISGAf and the other on TvISG (a combination of TvISGAf and TvISGAm), exhibited significant discrimination between positive and negative samples. ELISA based exclusively on TvISGAm demonstrated a significant overlap between the populations of positive and negative sera, as observed in the histogram (Fig. 4B). However, the inclusion of the TvISGAm antigen in the TvISG-based ELISA was improved when combined with TvISGAf. Therefore, we decided to evaluate the complete panel of sera using only the TvISGAf-based and TvISG-based ELISA.

**Figure 4:**
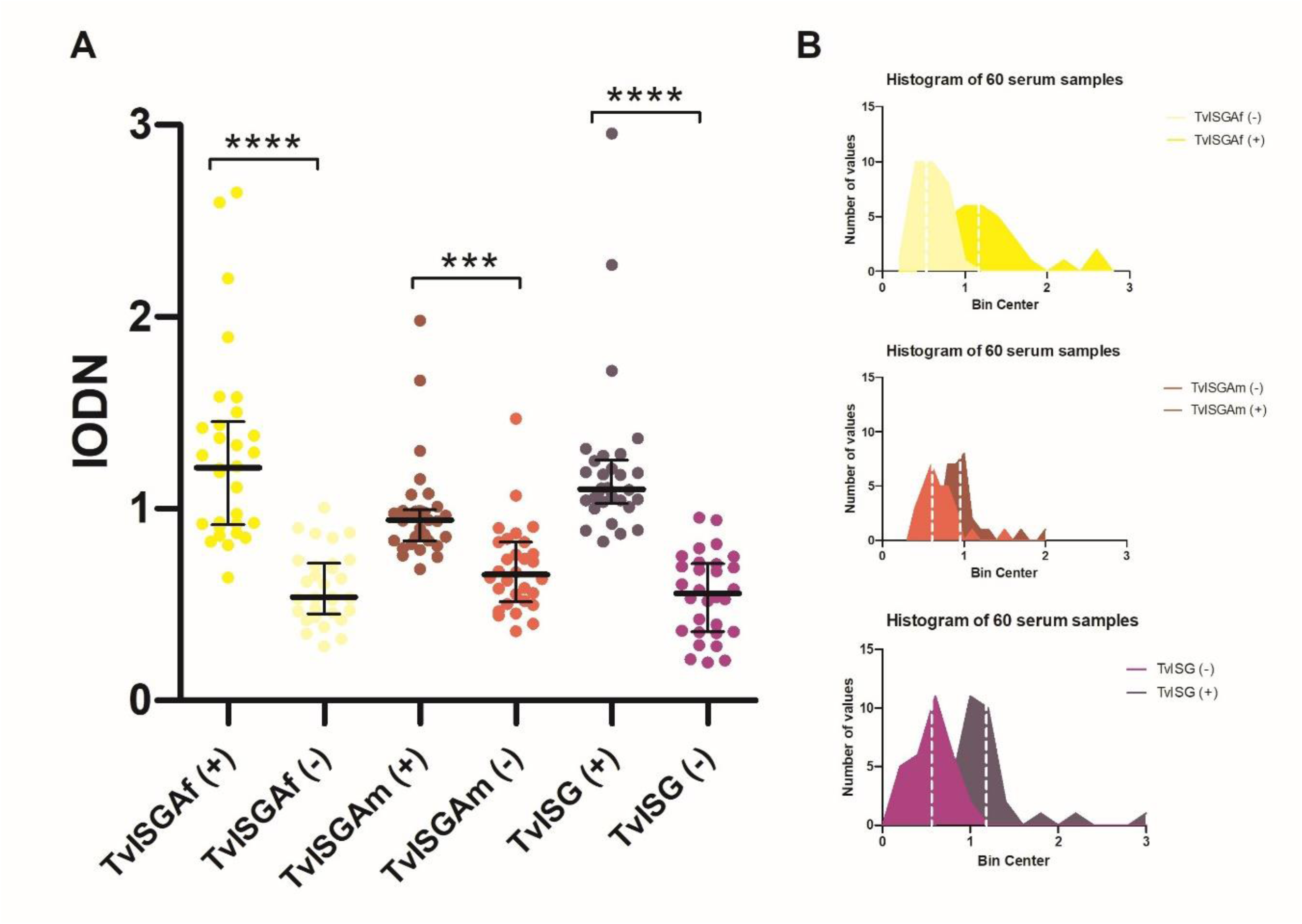
Comparing reactivity of TvISGAf (yellow) or TvISGAm (orange) antigens or a mixture of them (TvISG, purple) in ELISA using confirmed positive and negative results from dairy cattle sera. (A). Histograms indicate the distribution of the populations of positive and negative sera (B). *** *p* < 0.001, **** *p* < 0.0001.

Additionally, the antigens evaluated in the ELISA produced negative results (below the positivity threshold) when tested using bovine sera from animals infected with *Anaplasma marginale*, *Babesia bovis*, and *Trypanosoma theileri* (S7 Fig).

### Sensitivity and specificity of TvISGAf-based ELISA and TvISG-based ELISA

The formal parameters of sensitivity and specificity (proportion of correct negative results) for each test were determined by ROC curve analysis using 212 serum samples (Fig. 5). The combination of the two ISGs in the TvISG-based ELISA exhibited the most significant discrimination between *T. vivax*-infected bovines and control bovines, with an area under the ROC curve of 0.951, whereas that of the TvISGAf-based ELISA was 0.921. The TvISG-based ELISA utilizing the antigenic mixture achieved a sensitivity of 89.6% (95% CI of 82.4% to 93.2%) and a specificity of 93.8% (95% CI of 84.8% to 98.3%), whereas the sensitivity and specificity of the TvISGAf-based ELISA were 85.9% (95% CI of 79.1% to 91.1%) and 89.1% (95% CI of 78.8% to 95.5%), respectively. In addition, pairwise comparison of the ROC curves revealed significant differences between the mixture and TvISGAf (*p* = 0.0192).

**Figure 5:**
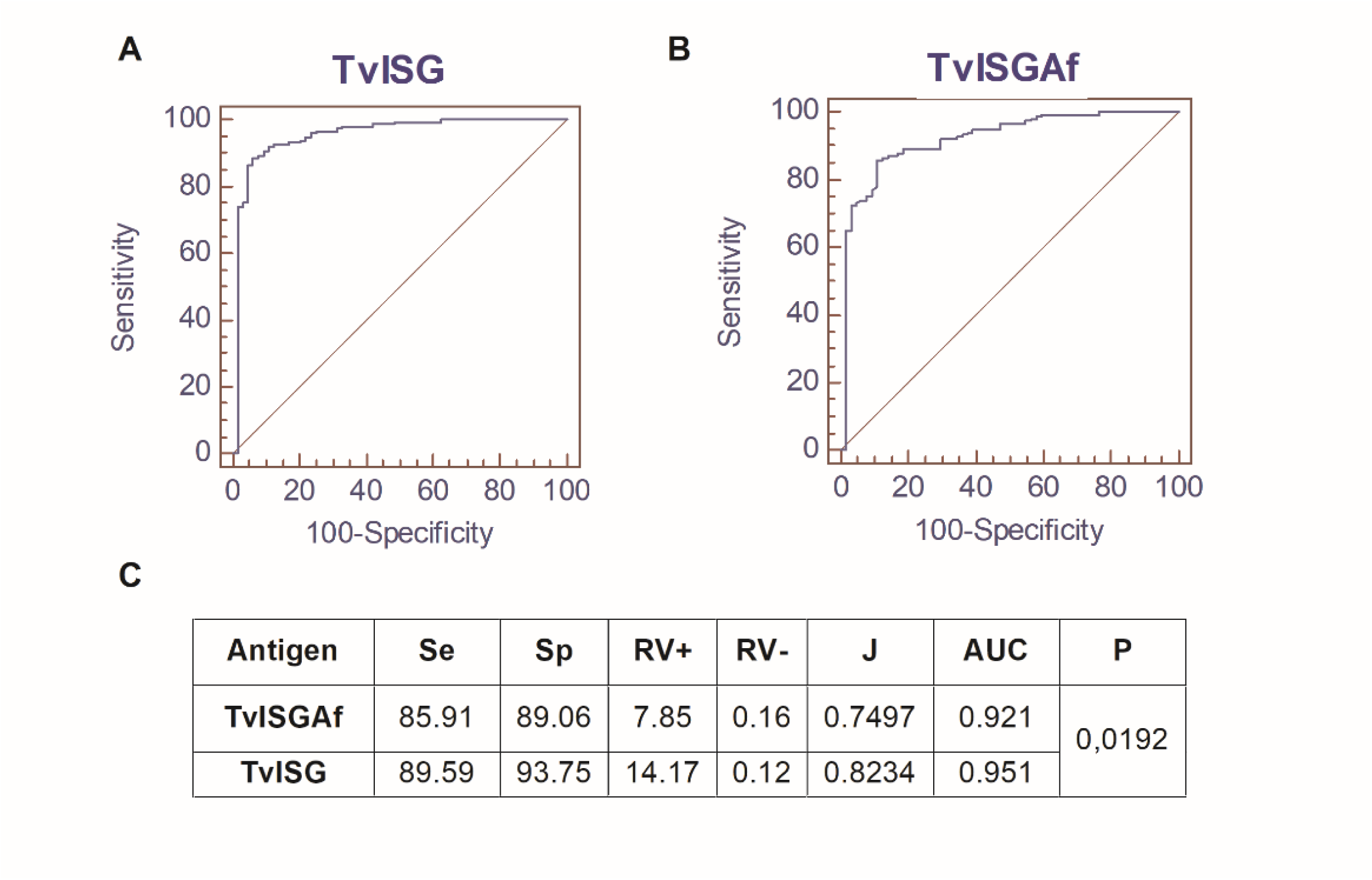
Comparison of the ROC curves of the anti-IgG ELISA tests of bovine sera using TvISGAf antigen and mixture TvISG. ROC curves for the TvISG mixture of TvISGAf and TvISGAm antigens (A) and TvISGAf (B). Different ELISA assays were performed using sera from the following groups: negative control (n = 63) and animals positive for *Trypanosoma vivax* in parasitological and PCR tests (n = 149). Table of diagnostic performance (C); Se, sensitivity; Sp, specificity; RV, likelihood ratio; J, Youden index; AUC, area under the ROC curve; P, pairwise comparison of ROC curves.

### Kinetics of detection using TvISG-based ELISA

The performance of the TvISG-based ELISA was assessed in sera from recently infected cattle (day 0), and at 15– and 30-days post-treatment with ISM. Two groups of cattle from two dairy farms were evaluated. One dairy farm recently reported cases of *T. vivax* infection (acute group), whereas the other dairy farm had chronic *T. vivax* infections, referred to as the chronic group (Fig. 6). Similar patterns were observed in the chronic group, with a slight increase in antibodies on day 15, followed by a decrease in the mean on day 30 (Pearson r = 0.5513; *p* = 0.6282). In the acute group, there was an evident increase in post-treatment IODN levels, indicating a correlation with time (Pearson r = 0.9986; *p* = 0.0341). This association exhibited an increasing trend, fitting into a linear model within the assessed and evaluated range (*R^2^* = 0.9971; *p* = 0.0341).

**Figure 6:**
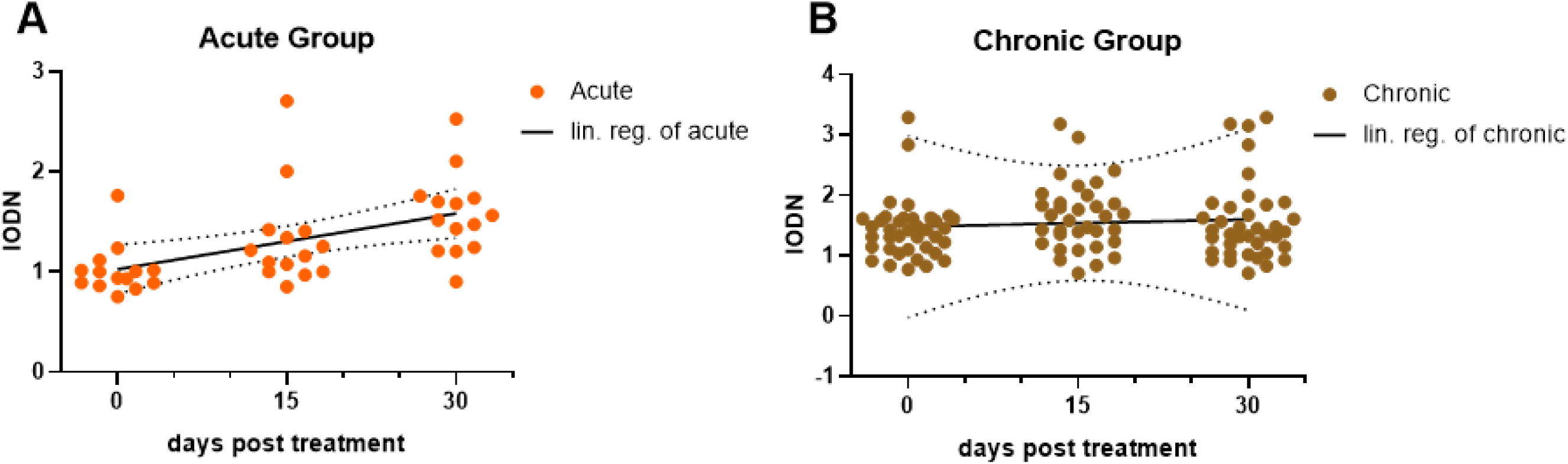
Kinetic Evaluation of Acute and Chronic Groups in Holstein Dairy Cattle. Positive serum samples were gathered from Holstein dairy cattle on day 0, following confirmation through parasitological and PCR tests. Subsequently, the cattle were administered Isometamidium. Follow-up samples from the same dairy cattle were obtained on days 15 and 30. Acute group consisted of 14 positive serum samples originating from a dairy farm with a recent report of *T. vivax* infection (A). Chronic group consisted of 36 positive serum samples were obtained from a dairy farm with a history of chronic *T. vivax* infection (B).

### Seroprevalence of bovine trypanosomiasis in the central region of Argentina as determined by TvISG-based ELISA

In this study, 892 samples collected from dairy cows in 40 fields in the central region of Argentina were screened using the TvISG-based ELISA (Fig. 1). Seroprevalence for bovine trypanosomiasis was assessed and is presented in Table 1. *T. vivax* infection was identified in 472 of 892 (53%) samples. Notably, the highest seroprevalence for *T. vivax* was observed in San Justo, Córdoba Province, and San Cristobal, Santa Fe Province, with rates of 69% and 68%, respectively. In other regions, seroprevalence in the different departments the Santa Fe province was 55% in San Martin, 50% in Castellanos, 42% in San Jeronimo, and 21% in Vera. Collectively, the province of Santa Fe had an overall seroprevalence rate of 47%.

**Table 1:**
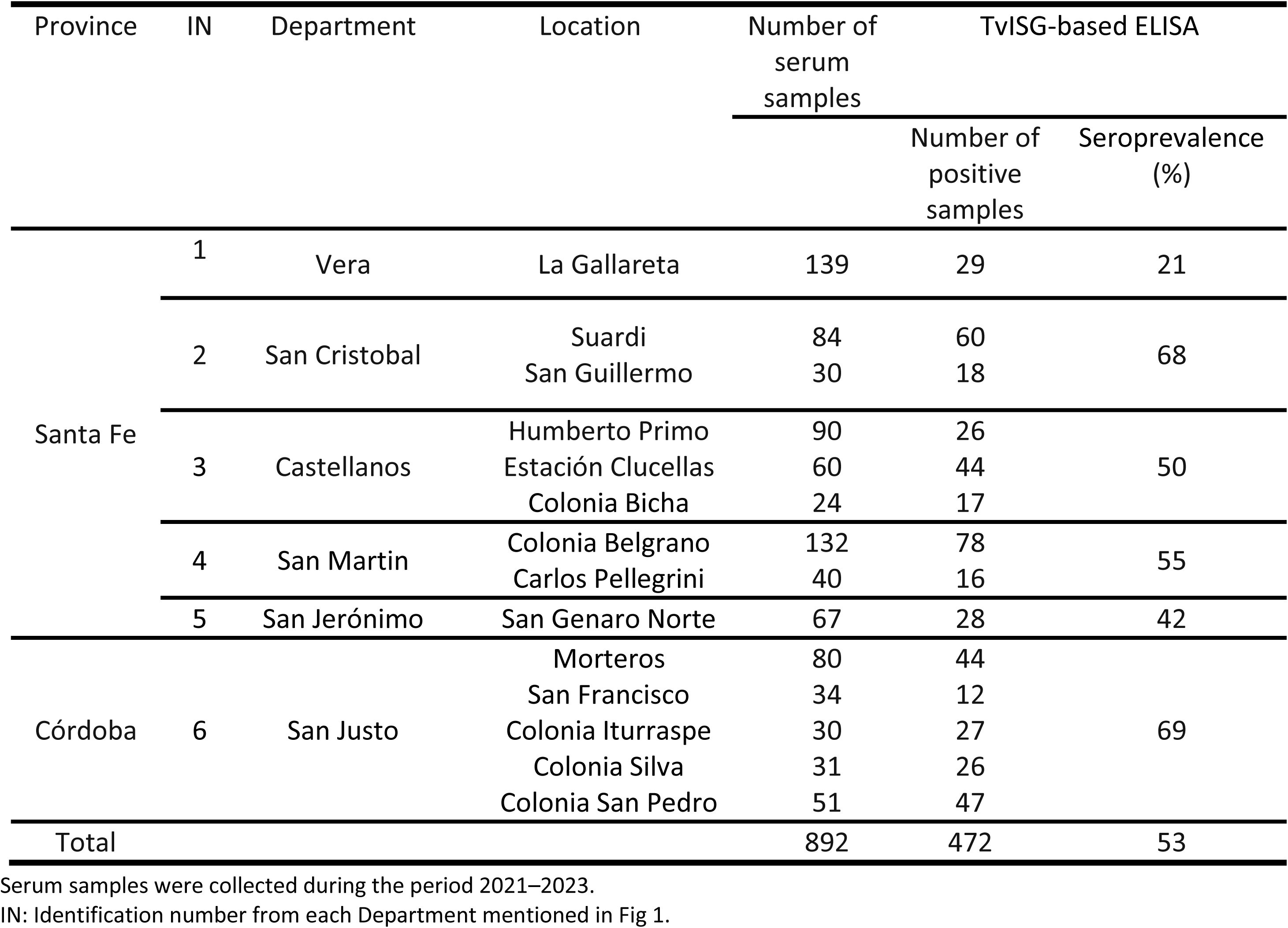
Seroprevalence of *T. vivax* among cattle (n = 892) in two provinces in Argentina was determined using TvISG-based ELISA.

## Discussion

Bovine trypanosomiasis caused by *T. vivax* is currently diagnosed using a combination of clinical, parasitological, molecular, and serological techniques (36–38). Parasitological diagnosis is the most commonly applied method for identifying *T. vivax* in the field, although its low sensitivity is only suitable for diagnosis during a short period of high parasitemia (39). IFAT and ELISA are the most commonly used diagnostic assays to determine the prevalence of trypanosomiasis. IFAT requires a host for parasite cultivation, making it less scalable and standardized (19). Crude extracts have been used to diagnose both *T. vivax* and *T. evansi* in the acute and chronic phases of the disease (40). Considering the cross-reactivity observed with extracts of both parasites (19,41), recombinant antigens have been proposed for specific serodiagnosis of *T. vivax* (6,36).

Various antigens have been proposed for the diagnosis of bovine trypanosomiasis (20–24). Fleming et al. (2016) identified two ISGs (TvY486_0019690 and TvY486_0045500) in African strain of *T. vivax*, with TvY486_0045500 showing the best diagnostic performance. In this study, we evaluated a truncated variant of antigen TvY486_0045500 (TvISGAf) against a panel of 212 serum samples from Argentina. Fleming et al. (2016) reported that TvY486_0045500 had sensitivity and specificity of 94.5% and 88.0%, respectively. However, our ELISA using the same antigen showed similar specificity but a 10% lower sensitivity. Although we used a smaller fraction, we omitted the non-globular region of the protein that was attached to the anchor site and did not present immunodominant B epitopes. However, this difference could be due to the fact that the sera evaluated by Fleming et al. (2016) came from African cattle, whereas the sera in our study came from Latin American cattle. In addition, the antigen cloned by Fleming et al. (2016) and reproduced by us as TvISGAf comes from the DNA of an African strain, Y468. Infecting strains in Latin America differ phylogenetically from African strains (42,43), which could contribute to the decrease in sensitivity. To enhance diagnostic performance, we evaluated an ISG homologue of TvY486_0045500 from a South American isolate (referred to as IB, (29)). TvISGAm shared approximately 63% identity and 69% similarity with TvISGAf. Furthermore, by assessing the localization of these ISGs, we demonstrated that the antibodies generated by the evaluated recombinant ISGs recognized the *T. vivax* membrane. (Figure 3). However, when we evaluated diagnostic capacity, TvISGAm did not yield favorable results. Nevertheless, the performance of this antigen combined with TvISGAf in detecting anti-*T. vivax* IgG in livestock serum demonstrated an increase in the Youden index by 0.074 and a statistically significant improvement in the area under the curve (p = 0.0192), indicating heightened sensitivity (over 3.7%) and specificity (over 4.7%) compared to the performance obtained for ELISA based on TvISGAf.

In this study, we continued our evaluation by using only the antigenic mixture, the TvISG-based Elisa, because of its superior diagnostic capability. We assessed the kinetics of antibody detection using the TvISG-based ELISA in cattle with acute and chronic *T. vivax* infections, as reported by dairy farms. In the chronic group, we observed robust antibody levels that persisted for up to 30 days post treatment with ISM. Notably, the acute group exhibited significant seropositivity from day 0 of treatment initiation, and antibody levels continued to increase 30 days after treatment. It is noteworthy that PCR results were negative at both the 15– and 30-day measurements; however, serological results indicated that this sustained increase in antibodies 30 days after treatment suggests parasite persistence in these individuals under the sensitivity of PCR. This aligns with the findings reported after ISM treatment of a naturally infected herd in Brazil (44).

Finally, we conducted a field seroprevalence study using TvISG-based ELISA in dairy animals (mostly females) in the central provinces of Argentina. We evaluated 40 establishments in our country’s dairy basin, focusing on the provinces of Santa Fe and Córdoba, which have been molecularly confirmed to have *T. vivax* infection. The overall seroprevalence was 53%, which aligns with the findings of Florentin et al. (2021), who molecularly tested approximately 150 samples and reported positivity rates ranging from 39% to 55% (17). The overall seroprevalence rate obtained in the province of Córdoba was nearly 20% higher than that in Santa Fe. However, numerous localities have been tested in Santa Fe province, with seroprevalence rates ranging from 21% to 68%. Argentina has thousands of dairy establishments, with nearly 95% of Argentine milk production coming from the provinces of Córdoba (37%), Santa Fe (32%), and Buenos Aires (25%), followed by Entre Ríos (50), however, to date Buenos Aires has no cases of *T. vivax*, so the assessment in the provinces of Cordoba and Santa Fe covers almost 70% of the dairy basin, where *T. vivax* is endemic (17).

Our results are comparable to those of other studies in South America, highlighting the high prevalence of this disease in endemic regions. An epidemiological study in the Pantanal region of Brazil using indirect ELISA reported an average seroprevalence rate of 56% (41). Moderate seroprevalence of *T. vivax* has been reported in other Latin American countries: 14% in Peru, 23% in Ecuador, 40% in Paraguay (45), and 27% in Bolivia (46). The 53% reported in Argentina indicates how the parasite continues to spread throughout South America, and given the lack of control and surveillance mechanisms, it may invade other countries outside Africa and Latin America (1), as seen recently in the case of Iran in Asia (47).

Although *T. vivax* was initially documented in Argentina in 2006 (15), it was not until the summers of 2016 and 2017 that the first outbreaks were recorded in dairy farms (48). It is important to note that the clinical signs of bovine trypanosomiasis caused by *T. vivax* are non-specific, rendering clinical diagnosis challenging and imprecise, particularly in the presence of other cattle diseases such as anaplasmosis, babesiosis, bovine leptospirosis, leukemia, neosporosis, and viral infections (14,37,49). The lack of awareness of the disease, the large number of registered dairy farms in the dairy area (50), the sale of infected animals, and the use of the same syringes for vaccination have contributed to its rapid spread between the provinces of Córdoba and Santa Fe. Significant economic impacts have been caused by losses in dairy cattle infected with *T. vivax*, leading to the treatment of entire herds and resulting in a substantial reduction in milk production from affected herds (17,48). In this study, we demonstrated high seroprevalence of the disease in an affected region, suggesting a possible spread of the disease in the southern and eastern areas of the Pampas region, as observed by other authors (17,51).

Control of bovine trypanosomiasis is of great concern in affected countries because of significant economic losses. Currently, no vaccine is available for this disease, and climate change favors the spread of vector insects to increasingly less temperate areas (52). Therefore, precise detection and control programs are necessary. The results of this study suggest the need for bovine trypanosomiasis surveillance programs not only in endemic regions, but also in other countries with intensive livestock production. The sensitivity and specificity of the TvISG-based ELISA, designed to detect specific antibodies against *T. vivax*, makes it a valuable test for inclusion in surveillance programs. In addition, given the need for point-of-care devices, the ISG antigen mixture cand be used to develop an immunochromatographic test, facilitating field testing.

## Ethical Approval

The study was conducted in strict accordance with the recommendations of the Argentinean National Council of Animal Experimentation and was approved by the Research Ethics and Safety Committee of the Facultad de Bioquimica y Ciencias Biologicas de la UNL (process number: CE2021-02). Handling and sampling of cattle were performed by trained personnel, with animal safety and welfare as priority, and in strict accordance with good animal practice guidelines. by the Animal. The authors confirm that the ethical policies of the journal, as noted on the journal’s author guidelines page, have adhered to.

## Supporting Information

**S1 Fig. Transmembrane domain prediction of TvY486_0045500 protein.** The transmembrane domain probability profile was generated using SOSUI (http://harrier.nagahama-i-bio.ac.jp/sosui/).

**S2 Fig. Transmembrane domain prediction of Invariant Surface Glycoprotein from American *T. vivax*.** The transmembrane domain probability profile was generated using SOSUI (http://harrier.nagahama-i-bio.ac.jp/sosui/).

**S3 Fig. Amino acid sequence alignment of Invariant Surface Glycoprotein from African and American *T. vivax*.** Amino acid alignment was constructed using the Clustal W algorithm.

**S4 Fig. Prediction of linear antigenic epitopes of Invariant Surface Glycoprotein from African and American *T. vivax*.** The prediction probability profile was generated using SVMTriP (http://sysbio.unl.edu/SVMTriP/index.php).

**S5 Fig. Full amino acid sequences of recombinant TvISGAf and TvISGAm proteins.** The sequences of the fusion-tags are highlighted in green, and the *T. vivax* proteins are indicated in yellow.

**S6 Fig. Negative control of immunodetection of ISG protein in *T. vivax* cells.** Fluorescence microscopy images were performed without primary antibodies (mouse anti-TvISGAf or mouse anti-TvISGAm or rabbit anti-cPx) in the presence of Alexa680 conjugated goat anti-IgG and FITC conjugated goat anti-IgG. DAPI was used for nuclear and kinetoplast staining (blue).

**S7 Fig. Evaluation of cross-reactivity in TvISG ELISA.** Cutoff values of the tests were 0.80 (indicated by broken lines). Samples from cattle infected with Babesia bovis (20), Anaplasma marginale (20), and Trypanosoma theileri (8).

**S1 Table: Primers used for amplifying the encoding sequences of Invariant Surface Glycoprotein from African and American *T. vivax*.**

## Reference

1. Fetene E, Leta S, Regassa F, Büscher P. Global distribution, host range and prevalence of Trypanosoma vivax: a systematic review and meta-analysis. Parasites & Vectors. 25 de enero de 2021;14(1):80.

2. Moloo SK, Sabwa CL, Kabata JM. Vector competence of Glossina pallidipes and G. morsitans centralis for Trypanosoma vivax, T. congolense and T. b. brucei. Acta Trop. agosto de 1992;51(3-4):271–80.

3. Seidl A, Dávila A, Silva R. Estimated financial impact of Trypanosoma vivax on the Brazilian pantanal and Bolivian lowlands. Memorias do Instituto Oswaldo Cruz [Internet]. abril de 1999 [citado 27 de febrero de 2023];94(2). Disponible en: https://pubmed.ncbi.nlm.nih.gov/10224541/

4. Cortez AP, Rodrigues AC, Garcia HA, Neves L, Batista JS, Bengaly Z, et al. Cathepsin L-like genes of Trypanosoma vivax from Africa and South America--characterization, relationships and diagnostic implications. Mol Cell Probes. febrero de 2009;23(1):44–51.

5. Gardiner PR. Recent studies of the biology of Trypanosoma vivax. Adv Parasitol. 1989;28:229–317.

6. Mekata H, Konnai S, Witola WH, Inoue N, Onuma M, Ohashi K. Molecular detection of trypanosomes in cattle in South America and genetic diversity of Trypanosoma evansi based on expression-site-associated gene 6. Infect Genet Evol. diciembre de 2009;9(6):1301–5.

7. Batista JS, Riet-Correa F, Teixeira MMG, Madruga CR, Simões SDV, Maia TF. Trypanosomiasis by Trypanosoma vivax in cattle in the Brazilian semiarid: Description of an outbreak and lesions in the nervous system. Vet Parasitol. 31 de enero de 2007;143(2):174–81.

8. Cadioli FA, Fidelis Junior OL, Sampaio PH, dos Santos GN, André MR, Castilho KJG de A, et al. Detection of Trypanosoma vivax using PCR and LAMP during aparasitemic periods. Vet Parasitol. 30 de noviembre de 2015;214(1-2):174–7.

9. Galiza GJN, Garcia HA, Assis ACO, Oliveira DM, Pimentel LA, Dantas AFM, et al. High mortality and lesions of the central nervous system in trypanosomosis by Trypanosoma vivax in Brazilian hair sheep. Vet Parasitol. 15 de diciembre de 2011;182(2-4):359–63.

10. Bastos TSA, Faria AM, Madrid DM de C, Bessa LC de, Linhares GFC, Fidelis Junior OL, et al. First outbreak and subsequent cases of Trypanosoma vivax in the state of Goiás, Brazil. Rev Bras Parasitol Vet. 29 de junio de 2017;26(3):366–71.

11. Jaimes-Dueñez J, Triana-Chávez O, Mejía-Jaramillo AM. Parasitological and molecular surveys reveal high rates of infection with vector-borne pathogens and clinical anemia signs associated with infection in cattle from two important livestock areas in Colombia. Ticks Tick Borne Dis. febrero de 2017;8(2):290–9.

12. Cadioli FA, Barnabé P de A, Machado RZ, Teixeira MCA, André MR, Sampaio PH, et al. First report of Trypanosoma vivax outbreak in dairy cattle in São Paulo state, Brazil. Rev Bras Parasitol Vet. junio de 2012;21(2):118–24.

13. Batista JS, Rodrigues CMF, Olinda RG, Silva TMF, Vale RG, Câmara ACL, et al. Highly debilitating natural Trypanosoma vivax infections in Brazilian calves: epidemiology, pathology, and probable transplacental transmission. Parasitol Res. enero de 2012;110(1):73–80.

14. Osório ALAR, Madruga CR, Desquesnes M, Soares CO, Ribeiro LRR, Costa SCG da. Trypanosoma (Duttonella) vivax: its biology, epidemiology, pathogenesis, and introduction in the New World--a review. Mem Inst Oswaldo Cruz. febrero de 2008;103(1):1–13.

15. Monzon CM, Mancebo OA, Giménez JN, Russo AM. Primera descripción de tripanosoma vivax en Argentina. septiembre de 2008 [citado 28 de febrero de 2023]; Disponible en: https://ri.conicet.gov.ar/handle/11336/60849

16. Paoletta MS, López Arias L, de la Fournière S, Guillemi EC, Luciani C, Sarmiento NF, et al. Epidemiology of Babesia, Anaplasma and Trypanosoma species using a new expanded reverse line blot hybridization assay. Ticks Tick Borne Dis. febrero de 2018;9(2):155–63.

17. Florentin AS, Garcia Perez HA, Rodrigues CMF., Dubois EF, Monzón CM, Teixeira MMG. Molecular epidemiological insights into Trypanosoma vivax in Argentina: From the endemic Gran Chaco to outbreaks in the Pampas. Transboundary and Emerging Diseases [Internet]. 21 de septiembre de 2021 [citado 30 de noviembre de 2021];n/a(n/a). Disponible en: https://onlinelibrary.wiley.com/doi/abs/10.1111/tbed.14103

18. Luckins AG, Mehlitz D. Evaluation of an indirect fluorescent antibody test, enzyme-linked immunosorbent assay and quantification of immunoglobulins in the diagnosis of bovine trypanosomiasis. Trop Anim Health Prod. agosto de 1978;10(3):149–59.

19. Luckins AG. Detection of antibodies in trypanosome-infected cattle by means of a microplate enzyme-linked immunosorbent assay. Trop Anim Health Prod. febrero de 1977;9(1):53–62.

20. Pinheiro GRG, Ferreira LL, Teixeira Silva AL, Cardoso MS, Ferreira-Júnior Á, Steindel M, et al. A recombinant protein (MyxoTLm) for the serological diagnosis of acute and chronic Trypanosoma vivax infection in cattle. Vet Parasitol. agosto de 2021;296:109495.

21. Boulangé A, Pillay D, Chevtzoff C, Biteau N, Comé de Graça V, Rempeters L, et al. Development of a rapid antibody test for point-of-care diagnosis of animal African trypanosomosis. Vet Parasitol. 15 de enero de 2017;233:32–8.

22. Pillay D, Izotte J, Fikru R, Büscher P, Mucache H, Neves L, et al. Trypanosoma vivax GM6 antigen: a candidate antigen for diagnosis of African animal trypanosomosis in cattle. PLoS One. 2013;8(10):e78565.

23. Camargo R, Izquier A, Uzcanga GL, Perrone T, Acosta-Serrano A, Carrasquel L, et al. Variant surface glycoproteins from Venezuelan trypanosome isolates are recognized by sera from animals infected with either Trypanosoma evansi or Trypanosoma vivax. Vet Parasitol. 15 de enero de 2015;207(1-2):17–33.

24. Fleming JR, Sastry L, Wall SJ, Sullivan L, Ferguson MAJ. Proteomic Identification of Immunodiagnostic Antigens for Trypanosoma vivax Infections in Cattle and Generation of a Proof-of-Concept Lateral Flow Test Diagnostic Device. PLOS Neglected Tropical Diseases. 8 de septiembre de 2016;10(9):e0004977.

25. Fleming JR, Sastry L, Crozier TWM, Napier GB, Sullivan L, Ferguson MAJ. Proteomic selection of immunodiagnostic antigens for Trypanosoma congolense. PLoS Negl Trop Dis. junio de 2014;8(6):e2936.

26. Ziegelbauer K, Multhaup G, Overath P. Molecular characterization of two invariant surface glycoproteins specific for the bloodstream stage of Trypanosoma brucei. J Biol Chem. 25 de mayo de 1992;267(15):10797–803.

27. Sullivan L, Wall SJ, Carrington M, Ferguson MAJ. Proteomic selection of immunodiagnostic antigens for human African trypanosomiasis and generation of a prototype lateral flow immunodiagnostic device. PLoS Negl Trop Dis. 2013;7(2):e2087.

28. Sambrook J, Russell DW. Molecular Cloning: A Laboratory Manual. Cold Spring Harbor, N.Y; 2000.

29. Greif G, Rodriguez M, Bontempi I, Robello C, Alvarez-Valin F. Different kinetoplast degradation patterns in American Trypanosoma vivax strains: Multiple independent origins or fast evolution? Genomics. 5 de enero de 2021;

30. Bradford MM. A rapid and sensitive method for the quantitation of microgram quantities of protein utilizing the principle of protein-dye binding. Anal Biochem. 7 de mayo de 1976;72:248–54.

31. Gonzalez LN, Cabeza MS, Robello C, Guerrero SA, Iglesias AA, Arias DG. Biochemical characterization of GAF domain of free-R-methionine sulfoxide reductase from Trypanosoma cruzi. Biochimie. 1 de octubre de 2023;213:190–204.

32. Bontempi I, Fleitas P, Poato A, Vicco M, Rodeles L, Prochetto E, et al. Trans-sialidase overcomes many antigens to be used as a vaccine candidate against Trypanosoma cruzi. Immunotherapy. junio de 2017;9(7):555–65.

33. Woo PT. The haematocrit centrifuge for the detection of trypanosomes in blood. Can J Zool. septiembre de 1969;47(5):921–3.

34. Boakye DA, Tang J, Truc P, Merriweather A, Unnasch TR. Identification of bloodmeals in haematophagous Diptera by cytochrome B heteroduplex analysis. Med Vet Entomol. julio de 1999;13(3):282–7.

35. Vicco MH, Bontempi IA, Rodeles L, Yodice A, Marcipar IS, Bottasso O. Decreased level of antibodies and cardiac involvement in patients with chronic Chagas heart disease vaccinated with BCG. Med Microbiol Immunol. abril de 2014;203(2):133–9.

36. Desquesnes M, Gonzatti M, Sazmand A, Thévenon S, Bossard G, Boulangé A, et al. A review on the diagnosis of animal trypanosomoses. Parasit Vectors. 19 de febrero de 2022;15(1):64.

37. Ferreira AVF, Garcia GC, Araújo FF de, Nogueira LM, Bittar JFF, Bittar ER, et al. Methods Applied to the Diagnosis of Cattle Trypanosoma vivax Infection: An Overview of the Current State of the Art. Current Pharmaceutical Biotechnology. 22 de septiembre de 2022;24(3):355–65.

38. Gonzatti MI, González-Baradat B, Aso PM, Reyna-Bello A. Trypanosoma (Duttonella) vivax and Typanosomosis in Latin America: Secadera/Huequera/Cacho Hueco. En: Magez S, Radwanska M, editores. Trypanosomes and Trypanosomiasis [Internet]. Vienna: Springer; 2014 [citado 13 de septiembre de 2023]. p. 261–85. Disponible en: 10.1007/978-3-7091-1556-5_11

39. Masake RA, Majiwa PA, Moloo SK, Makau JM, Njuguna JT, Maina M, et al. Sensitive and specific detection of Trypanosoma vivax using the polymerase chain reaction. Exp Parasitol. febrero de 1997;85(2):193–205.

40. Ramírez-Iglesias JR, Eleizalde MC, Gómez-Piñeres E, Mendoza M. Trypanosoma evansi: a comparative study of four diagnostic techniques for trypanosomosis using rabbit as an experimental model. Exp Parasitol. mayo de 2011;128(1):91–6.

41. Madruga CR, Araújo FR, Cavalcante-Goes G, Martins C, Pfeifer IB, Ribeiro LR, et al. The development of an enzyme-linked immunosorbent assay for Trypanosoma vivax antibodies and its use in epidemiological surveys. Mem Inst Oswaldo Cruz. noviembre de 2006;101(7):801–7.

42. Cortez AP, Ventura RM, Rodrigues AC, Batista JS, Paiva F, Añez N, et al. The taxonomic and phylogenetic relationships of Trypanosoma vivax from South America and Africa. Parasitology. agosto de 2006;133(Pt 2):159–69.

43. Greif G, Ponce de Leon M, Lamolle G, Rodriguez M, Piñeyro D, Tavares-Marques LM, et al. Transcriptome analysis of the bloodstream stage from the parasite Trypanosoma vivax. BMC Genomics. 5 de marzo de 2013;14:149.

44. Castilho Neto KJG de A, Garcia AB da CF, Fidelis Junior OL, Nagata WB, André MR, Teixeira MMG, et al. Follow-up of dairy cattle naturally infected by Trypanosoma vivax after treatment with isometamidium chloride. Rev Bras Parasitol Vet. 26 de abril de 2021;30:e020220.

45. Silva RA, Egüez A, Morales G, Eulert E, Montenegro A, Ybañez R, et al. Bovine trypanosomiasis in Bolivian and Brazilian lowlands. Mem Inst Oswaldo Cruz. 1998;93(1):29–32.

46. Gonzales JL, Loza A, Chacon E. Sensitivity of different Trypanosoma vivax specific primers for the diagnosis of livestock trypanosomosis using different DNA extraction methods. Vet Parasitol. 15 de marzo de 2006;136(2):119–26.

47. Asghari MM, Rassouli M. First identification of Trypanosoma vivax among camels (Camelus dromedarius) in Yazd, central Iran, jointly with Trypanosoma evansi. Parasitol Int. febrero de 2022;86:102450.

48. Abdala AA, Larriestra AJ, Signorini M, Abdala AA, Larriestra AJ, Signorini M. Estimación de pérdidas económicas causadas por Trypanosoma vivax en un rodeo lechero de Argentina. Revista veterinaria. julio de 2020;31(2):115–9.

49. Desquesnes M. Livestock trypanosomoses and their vectors in Latin America. Paris: OIE; 2004. 174 p.

50. SENASA. Tambos [Internet]. 2015 [citado 13 de septiembre de 2023]. Disponible en: https://www.senasa.gob.ar/prensa/DNSA/Control_Gestion_y_Programas_Especiales/Indicadores_ganaderos/7_Indicadores_Ganaderia_Bovina_%20de_Tambo/Tambos.html

51. Monzon CM. Evolución de la Tripanosomosis bovina po Trypanosoma vivax en Formosa (Argentina) Años 2007-2012 y su potencial dispersión en el país. 2013;

52. Mojahed N, Mohammadkhani MA, Mohamadkhani A. Climate Crises and Developing Vector-Borne Diseases: A Narrative Review. Iran J Public Health. diciembre de 2022;51(12):2664–73.

